# Sunday driver mediates multi-compartment Golgi outposts defects induced by amyloid precursor protein

**DOI:** 10.1101/2020.09.02.280321

**Authors:** Qianqian Du, Jin Chang, Guo Cheng, Yinyin Zhao, Wei Zhou

## Abstract

Golgi defects including Golgi fragmentation are pathological features of Alzheimer’ disease (AD). As a pathogenic factor of AD, amyloid precursor protein (APP) induces Golgi fragmentation in soma. However, how APP regulates Golgi outposts (GOs) in dendrites remains unclear. Given that APP resided and affected GOs movements, especially reversed the distribution of multi-compartment GOs (mcGOs), we investigated the regulatory mechanism of mcGOs movements in *Drosophila* larvae. Knockdown experiments showed the bidirectional mcGOs movements were cooperatively controlled by dynein heavy chain (Dhc) and kinesin heavy chain subunits. Notably, only Dhc mediated APP’s regulation on mcGOs movements. Further, by loss-of-function screening, the adaptor protein Sunday driver (Syd) was identified to mediate APP-induced alteration of the direction of mcGOs movements, and dendritic defects. Collectively, by elucidating a model of bidirectional mcGOs movements, we revealed the mechanism of APP’s regulation on the direction of mcGOs movements. It provides new insights into AD pathogenesis.

## Introduction

The Golgi apparatus, an organelle with unique cisternal stacked structure, is responsible for protein trafficking, sorting and processing. Accumulated evidences indicate that there are defects of the structure and function of the Golgi apparatus in AD brains. For example, fragmentation and atrophy of the Golgi apparatus has been observed in AD neurons (S. J. Baloyannis, 2014; Stavros J. Baloyannis & Ma, 2000; Stieber, Mourelatos, & Gonatas, 1996). Abnormalities of protein glycosylation also have been reported in AD brain tissues, such as an enhancement of glycosylation of the Tyr residue of Aβ peptides (Halim et al., 2011), and a decrease of neural sialyltransferase activity (Maguire & Breen, 1995). These results suggest that defects of the Golgi apparatus are pathological features of AD.

Previous studies suggest that defects of the Golgi apparatus are induced by amyloid precursor protein (APP). APP, the precusor of β-amyloid (Aβ), is one of the key pathogenic factors of AD (Golde, Estus, Younkin, Selkoe, & Younkin, 1992; Hardy & Higgins, 1992; Sisodia & Price, 1995). Studies have shown that overexpressing APP in neurons leads to Golgi fragmentation, companied by swollen cisternae and disorganized stacks (Joshi, Chi, Huang, & Wang, 2014). And, the mechanism of Golgi fragmentation is elucidated: the accumulation of Aβ produced by amyloidogenic APP processing induces phosphorylation of Golgi structural proteins, such as GRASP65 (Joshi et al., 2014) and GM130 (Sun et al., 2008). Hence, a hypothesis is raised: overexpression of APP leads to Golgi defects, then induces degeneration of the structure and function of neurons, thereby accelerating AD development.

In neurons, the Golgi apparatus exists not only in the soma but also in the dendrites, called Golgi outposts (GOs) (Horton & Ehlers, 2003; Horton et al., 2005; Pierce, Mayer, & Mccarthy, 2001). Compared with somal Golgi apparatus, dendritic GOs have their own unique features in the structure and function. GOs have different compartmental organization, including single-compartment GOs (scGOs) and multi-compartment GOs (mcGOs), with the disconnected *cis-*, *medial-*, and *trans-* cisternae in the former while stacked in the latter (Zhou et al., 2014). Meanwhile, different from the somal Golgi apparatus with stationary positioning, GOs are highly mobile, travelling anterogradely (towards dendritic ends) or retrogradely (towards the soma) (Horton & Ehlers, 2003; Ye et al., 2007). The dynamic GOs are involved in local processing and transport of proteins (Horton & Ehlers, 2003; Valenzuela & Perez, 2015), for example the NMDA receptors (NMDARs) are selectively transported to synapses via GOs while bypassing the Golgi apparatus (Jeyifous et al., 2009). And, the stationary GOs also have been found to be involved in the growth and stability of dendritic branches by functioning as local acentrosomal microtubule nucleation sites (Ori-McKenney, Jan, & Jan, 2012). Alterations of GOs have been reported in neurodegenerative diseases, for example, a loss of GOs in models of Machado-Joseph’s disease (Chung et al., 2017) and amyotrophic lateral sclerosis (Park et al., 2020), and abnormal GOs dynamic induced by leucine-rich repeat kinase (Lrrk), a Parkinson’s disease associated protein (Lin et al., 2015). However, the regulation of APP on GOs remains unknown.

Here, inspired by the finding that APP altered the distribution of mcGOs, we investigated the molecular mechanism of APP regulating the direction of mcGOs movements. First, *in vivo* imaging exhibited the correlation between APP and GOs through their colocalization, and the dynamic and distribution of GOs. Then, by knockdown of the motor proteins subunits, we investigated the mechanism that determined the direction of mcGOs movements. Further, the subunit which mediated alteration of the direction of mcGOs movements induced by APP was determined. Finally, by loss-of-function screening, the adaptor protein was discovered which was specifically involved in APP’s regulation on the direction of mcGOs movements, and its rescue on dendritic morphology was explored.

## Results

### APP alters the distribution of mcGOs in the dendrites

To investigate the regulation of APP on GOs, we first examined the correlation between APP and GOs by colocalization analysis. Using fluorescent protein-tagged APP (APP-GFP) and GalT (GalT-TagRFPt, a *trans*-Golgi marker), we observed the location of APP and GOs in dendrites of C3da neurons, a classic model for dendrite development (Grueber, Jan, & Jan, 2002). APP presented as punctate patterns in both soma and dendrites, and colocalized with somal Golgi apparatus and dendritic GOs (Fig 1a). Quantification of the overlapped GalT/APP showed that it accounted for about 84.6% of GalT (Fig 1b) and 49.6% of APP in the dendrites (Fig 1c). Similar results were also obtained between APP and ManII (ManII-TagRFPt, a *medial*-Golgi marker) (Figs 1a-c). Meanwhile, *in vivo* time-lapse imaging showed that both GOs and APP were dynamic in the dendrites (Fig 1d). Notably, the proportion of mobile GOs was decreased in APP-colocalized GOs when compared to GOs-alone puncta (Fig 1e). Besides, it was not changed between GOs-alone and GOs in wild-type neurons (Fig 1e). These results indicate that APP resides in the vast majority of GOs and alters the movements of GOs.

**Fig 1.**
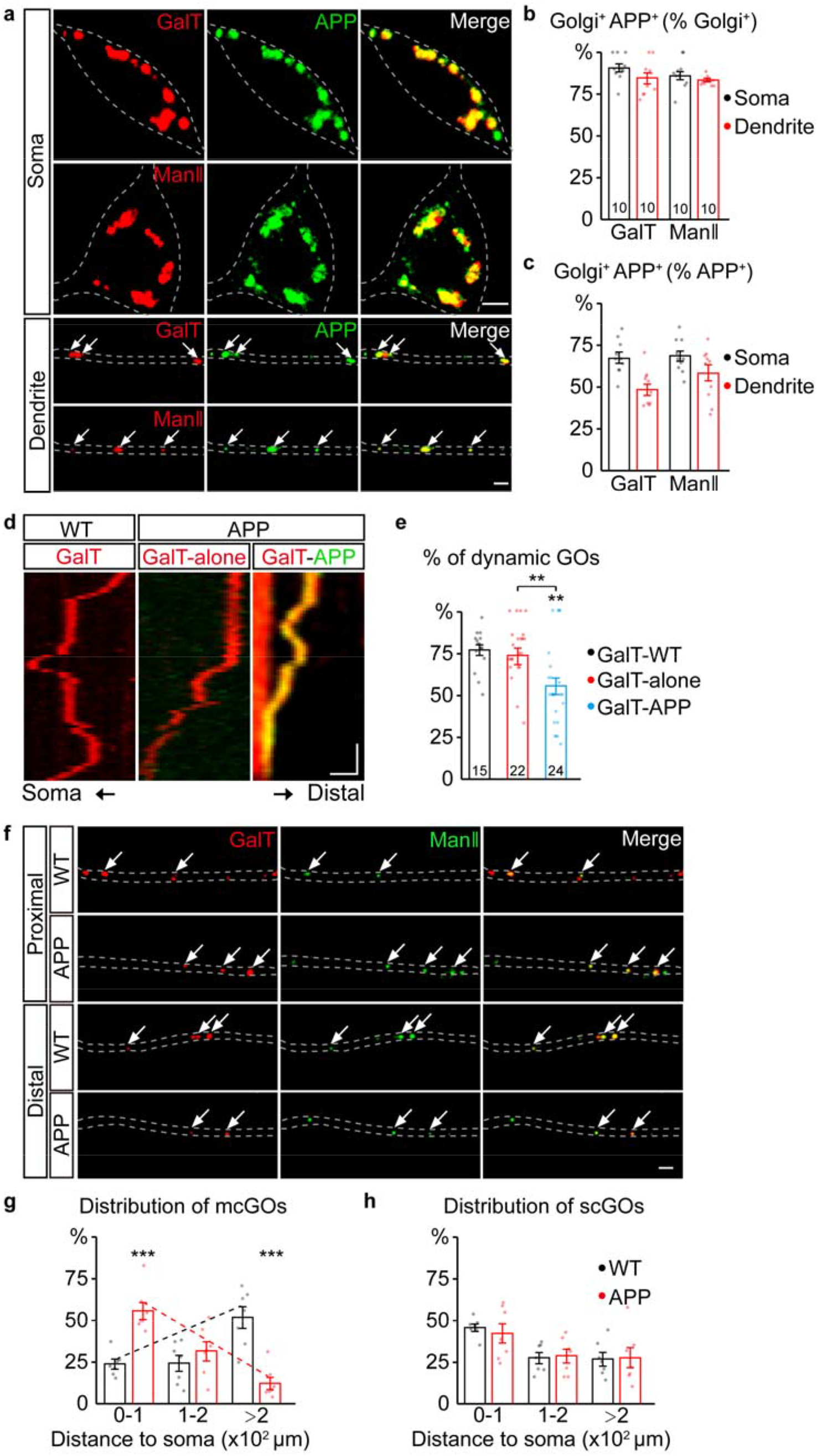
APP reverses the distribution of multi-compartment Golgi outposts (mcGOs) in the dendrites. **(a-c)** APP residing in the vast majority of GOs. **(a)** Examples of APP (green) and Golgi complex (red) in the soma and dendrites. APP is labeled by APP-GFP and Golgi complex is labeled by GalT-TagRFPt (top, *trans*-Golgi) or ManII-TagRFPt (bottom, *medial*-Golgi). Arrows indicate colocalized APP and GOs in the dendrites. **(b-c)** Quantification of the proportion of colocalized APP and Golgi complex in soma and dendrites, colocalization as a percentage of Golgi complex (**b**) or APP (**c**). **(d-e)** The residing APP changes the dynamic of GOs. **(d)** Examples of dynamic GOs with APP or not. Kymograph of time-lapse imaging of GOs showing GOs (labeled by GalT-TagRFPt) dynamic without (red) or with APP-GFP (yellow, GalT-APP). Scale bar: 2 μm / 1min. **(e)** Bar charts showing the proportion of dynamic GOs with APP or not. **(f-h)** APP reverses the distribution of mcGOs in the dendrites rather than single-compartment outposts (scGOs). **(f)** Examples of the distribution of GOs in wild-type and APP neurons. The *medial*- and *trans*-Golgi are labeled by ManII-GFP (green) and GalT-TagRFPt (red), respectively. Arrows point to mcGOs. **(g-h)** the distribution patterns of **(g)** mcGOs, and **(h)** scGOs in dendrites. Scale bar: 2 μm.

We next examined the distribution of GOs in dendrites (Fig 1f). Considering the existence of both stacked mcGOs and disconnected scGOs in the dendrites (Zhou et al., 2014), the fluorescently tagged *medial*- and *trans*-Golgi markers (ManII-GFP and GalT-TagRFPt) were used simultaneously. To quantify the distribution of GOs, the dendrites were divided into three segments according to the distance to soma: proximal (0 −100 μm), medial (100 - 200 μm), and distal (> 200 μm). The results showed that the distribution of mcGOs had a pronounced shift between wild-type and neurons which overexpressed human APP with Swedish mutation (APP neurons): the mcGOs were enriched in the proximal part of dendrite in APP neurons, whereas they dominated in the distal part in wild-type (Fig 1g). However, the distribution of scGOs showed no change in APP neurons (Fig 1h). These indicate that APP shifts the distribution tendency of mcGOs, from being enriched at distal dendrite to proximal dendrite.

In summary, we found that the overwhelming majority of GOs were resided by APP, and this residing altered GOs movements and reversed the distribution of mcGOs.

### Dhc and Khc cooperatively maintain the equilibrium state of the direction of mcGOs movements

Given that the distribution of mcGOs is related to the direction of their movements, the driving mechanism of mcGOs movements was studied. Here, we utilized a pipeline to show the trajectories of GOs movements (Figure 2a). It included three steps: time-lapse imaging, straightening of dendrites, generation and analysis of the kymograph of GOs movements. *In vivo* imaging showed that both mcGOs and scGOs can move bi-directionally in the dendrites (Figs 2a and b): anterograde and retrograde. Motor proteins, including kinesin and dynein, have been reported to be involved in GOs transport (Kelliher et al., 2018; Zheng et al., 2008). To find out the motor proteins which controlled mcGOs dynamic, 10 candidate subunits of motor proteins were tested using RNA-interference (RNAi) (Table S1). Results found that three RNAi lines led to a decrease of the proportion of dynamic GOs, which came from *Dhc (Dhc-64c*, *dynein heavy chain subunit)* and *Khc* (*kinesin heavy chain subunit*), as well as *Klc* (*kinesin light chain subunit*) (Fig S1a). Among them, *Dhc* and *Khc* were found to regulate the direction of dynamic GOs (Figs S1b and 2c). Loss of *Dhc* and *Khc* had opposite effects on the direction of mcGOs movements, with *Dhc* RNAi decreasing the proportion of anterograde movements of mcGOs, while *Khc* RNAi increasing it (Fig 2d). These indicate that *Dhc* is responsible for anterograde movements of mcGOs, while *Khc* mediates retrograde movements. Besides, the results also found that the direction of scGOs movements was only regulated by *Khc* RNAi rather than *Dhc* RNAi (Fig 2e). Neither *Dhc* RNAi nor *Khc* RNAi affected the displacements of mcGOs and scGOs movements (Figs 2f and g). Taken together, these results raise a model for the bidirectional movements of mcGOs: the anterograde and retrograde movements of mcGOs are drove by *Dhc* and *Khc* respectively, and the cooperation of *Dhc* and *Khc* maintains the equilibrium of the direction of mcGOs movements.

**Fig 2.**
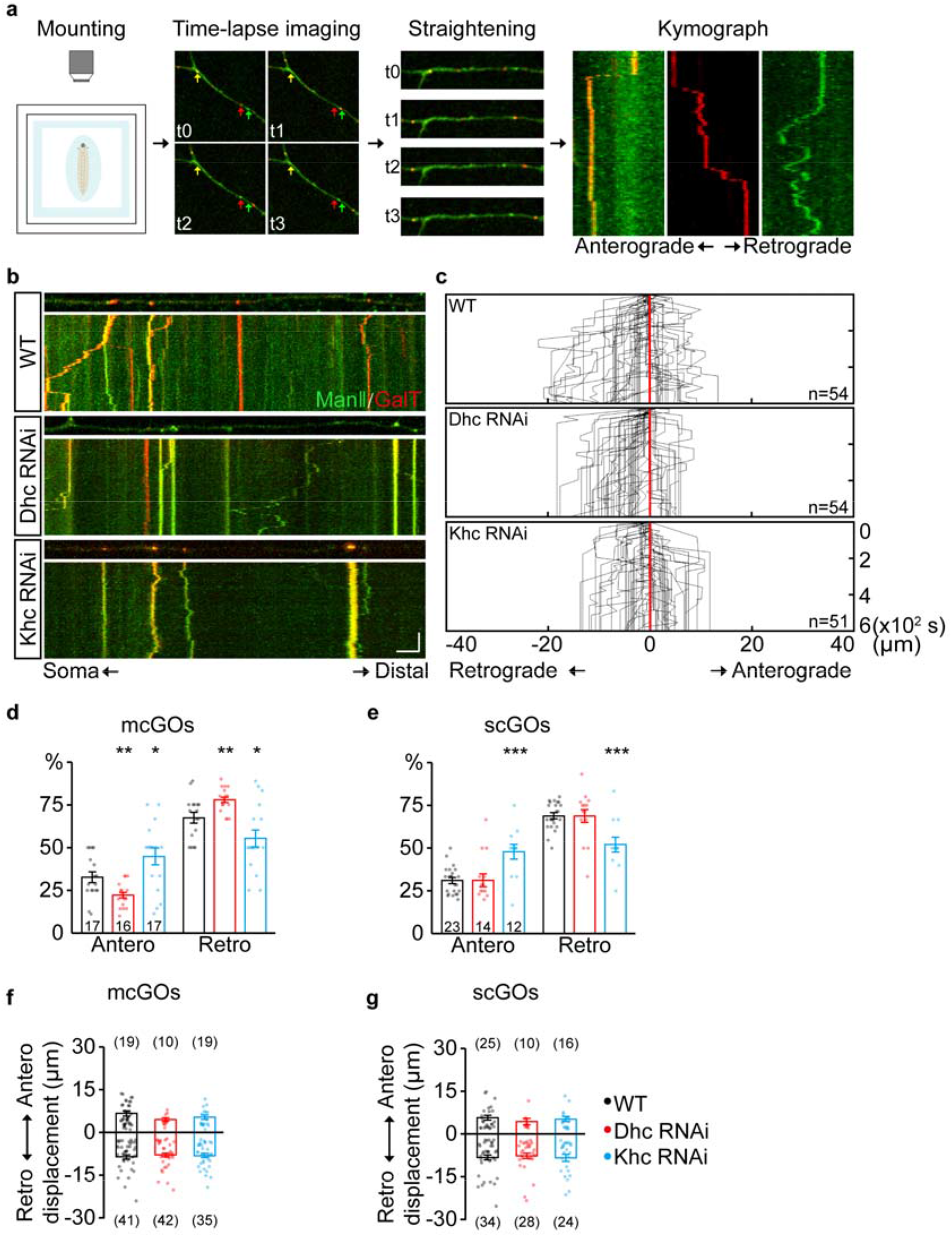
Dhc and Khc coordinate to maintain the equilibrium state of anterograde/retrograde movements of mcGOs. **(a)** Schematic diagram shows the pipeline for the generation of GOs trajectories. The four steps are (from left to right): the anesthetic larvae were 1) mounted for subsequent 2) time-lapse imaging. Arrows with different colors in the image indicate GOs with different organization. 3) the dynamic puncta in the dendrites of consecutive images were tracked down by the segmented line tool and 4) processed to generated a kymograph. **(b)** Examples of the trajectories of GOs dynamic in wild-type and APP neurons with RNAi knockdown of dynein heavy chain (*Dhc*) or kinesin heavy chain (*Khc*). Top: snapshots of the GOs in the straightened dendrites. Bottom: kymograph of time-lapse imaging of GOs in 10 min. Dual-colors live imaging showing the compartmental organization of GOs in dendrites, which are labeled by ManII-GFP (green, *medial*-Golgi) and GalT-TagRFPt (red, *trans*-Golgi). Green or red indicates a scGO and yellow shows mcGO. Scale bar: 5 μm / 2 min. **(c)** Compound tracks of dynamic GOs. Red lines are aligned along with displacement 0; puncta numbers are indicated within the boxes. **(d-e)** Quantification of the percentage of anterograde and retrograde movements for **(d)** mcGOs, **(e)** scGOs. The percentage of anterograde (retrograde) movements is calculated by dividing the number of anterograde (retrograde) moving mcGOs by the total number of moving mcGOs. **(f-g)** Quantification of the displacements of (**f**) mcGOs and (g) scGOs. Anterograde displacements are shown as positive, and retrograde as negative.

### APP enhances the anterograde movements of mcGOs driven by Dhc

Further, the dynamic behaviors of GOs in APP neurons were analyzed. APP also led to a decrease of the percentage of dynamic GOs (Figs 3a and b), similar to knockdown of *Dhc* and *Khc*. The anterograde and retrograde movements of GOs in APP neurons were quantified (Fig 3c). Results showed that the anterograde movements of mcGOs were increased in APP neurons (Fig 3d). Meanwhile, the displacements of retrograde movements of mcGOs in APP neurons became shorter when compared to wild-type, although the displacements of anterograde movements remained unchanged (Fig 3e). Taken together, these results indicate that APP alters the direction and displacements of mcGOs movements. In addition, the results also showed that the dynamic behaviors of scGOs in APP neurons had similar changes as mcGOs (Figs 3f and g).

**Fig 3.**
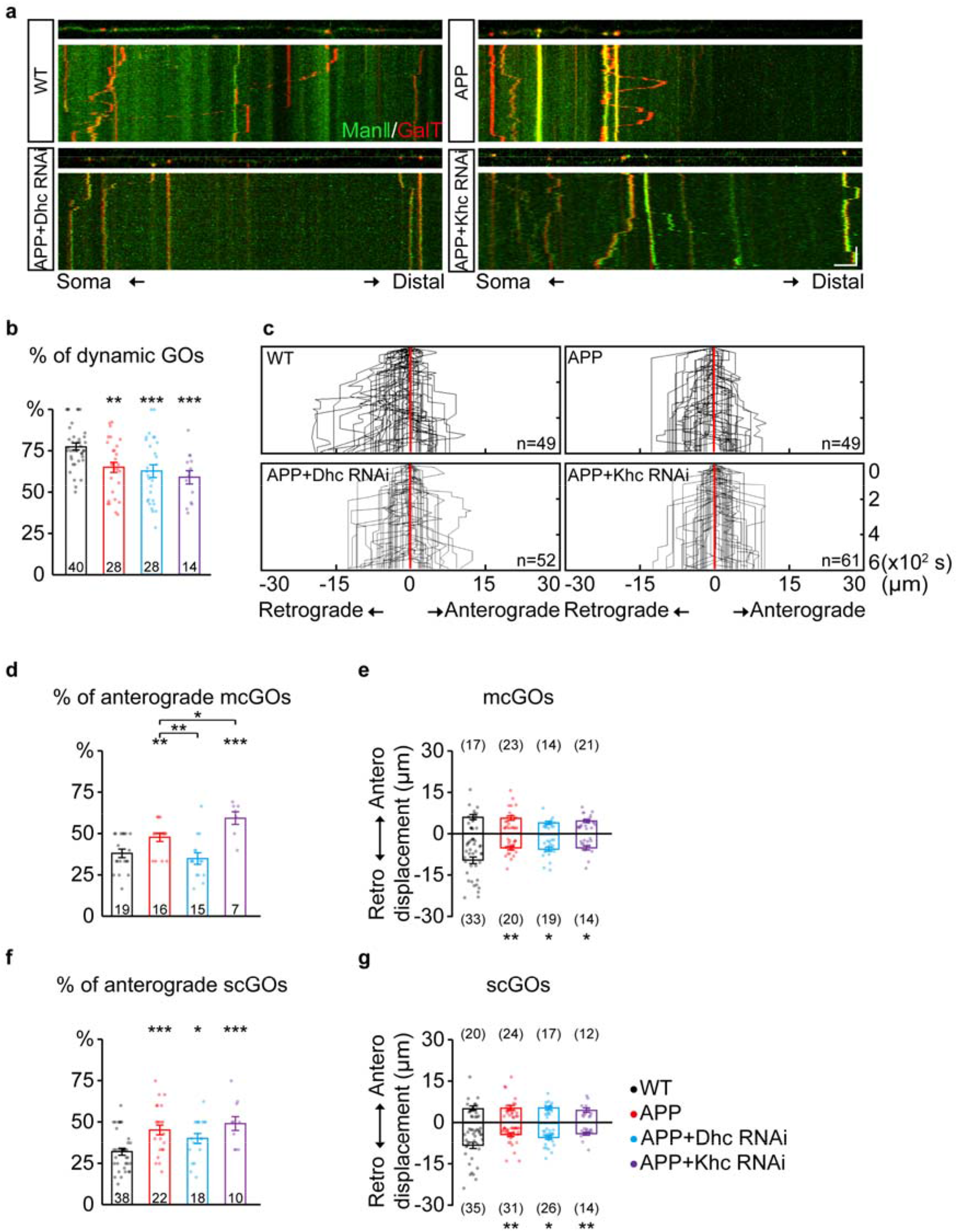
APP enhances the anterograde movements of mcGOs driven by *Dhc*. **(a)** Examples of the trajectories of GOs dynamic in wild-type and APP neurons with RNAi knockdown of dynein heavy chain (*Dhc*) or kinesin heavy chain (*Khc*). Top: snapshots of the GOs in the straightened dendrites. Bottom: kymograph of time-lapse imaging of GOs in 10 min. Scale bar: 5 μm / 2 min. **(b)** Quantification of the proportion of dynamic GOs. **(c)** Compound tracks of dynamic GOs. **(d-e)** Quantification of the features of mcGOs dynamic. **(d)** the percentage of anterograde movements, **(e)** displacements. **(f-g)** Quantification of the features of scGOs dynamic. **(f)** the percentage of anterograde movements. **(g)** displacements.

Further, to explore the contributions of *Dhc* and *Khc* to APP-induced alterations of GOs movements, we examined the dynamic behaviors of GOs in APP neurons after the knockdown of *Dhc* and *Khc*. The results showed that the proportion of anterograde movements of mcGOs only had no difference in APP neurons with *Dhc* RNAi rather than with *Khc* RNAi, compared with wild-type neurons (Fig 3d). This demonstrated that loss of *Dhc*, but not *Khc*, rescued the abnormal anterograde movements of mcGOs in APP neurons. While, they did not restore the displacements of mcGOs movements (Fig 3e). Besides, neither *Dhc* RNAi nor *Khc* RNAi can recover the alterations of scGOs movements induced by APP (Figs 3f and g).

Therefore, we discovered that the anterograde movements of mcGOs were enhanced in APP neurons, and this alteration can only be recovered via the loss of *Dhc* but not *Khc*.

### Sunday driver (Syd) specifically mediates the abnormality of the direction of mcGOs movement induced by APP

The adaptor proteins act as bridges for motor proteins to recognize and anchor specific cargos (Karcher, Deacon, & Gelfand, 2002). To identify the adaptor proteins that associated with mcGOs, we performed a loss-of-function screening for 24 adaptor proteins that had been reported to be involved in cargos transport (Table S2 and Fig 4a). By comparing the motility of GOs, we found 5 proteins, which were Htt, Lva, Miro, NudE and Syd: knockdown of them induced a decrease of the proportion of dynamic GOs (Fig S2a). Next, we examined the direction of GOs movements. Only Syd, Lva and NudE played a role in it (Fig S2b, Figs 4b and d). Syd was specific to mcGOs: *Syd* RNAi led to a decrease of the proportion of anterograde movements of mcGOs, but not scGOs (Figs 4e and f). Lva and NudE regulated both mcGOs and scGOs. The difference between the function of Lva and NudE was that Lva^DN^ decreased the proportion of anterograde movements, while *NudE* RNAi increased it (Figs S3a-c).

**Fig 4.**
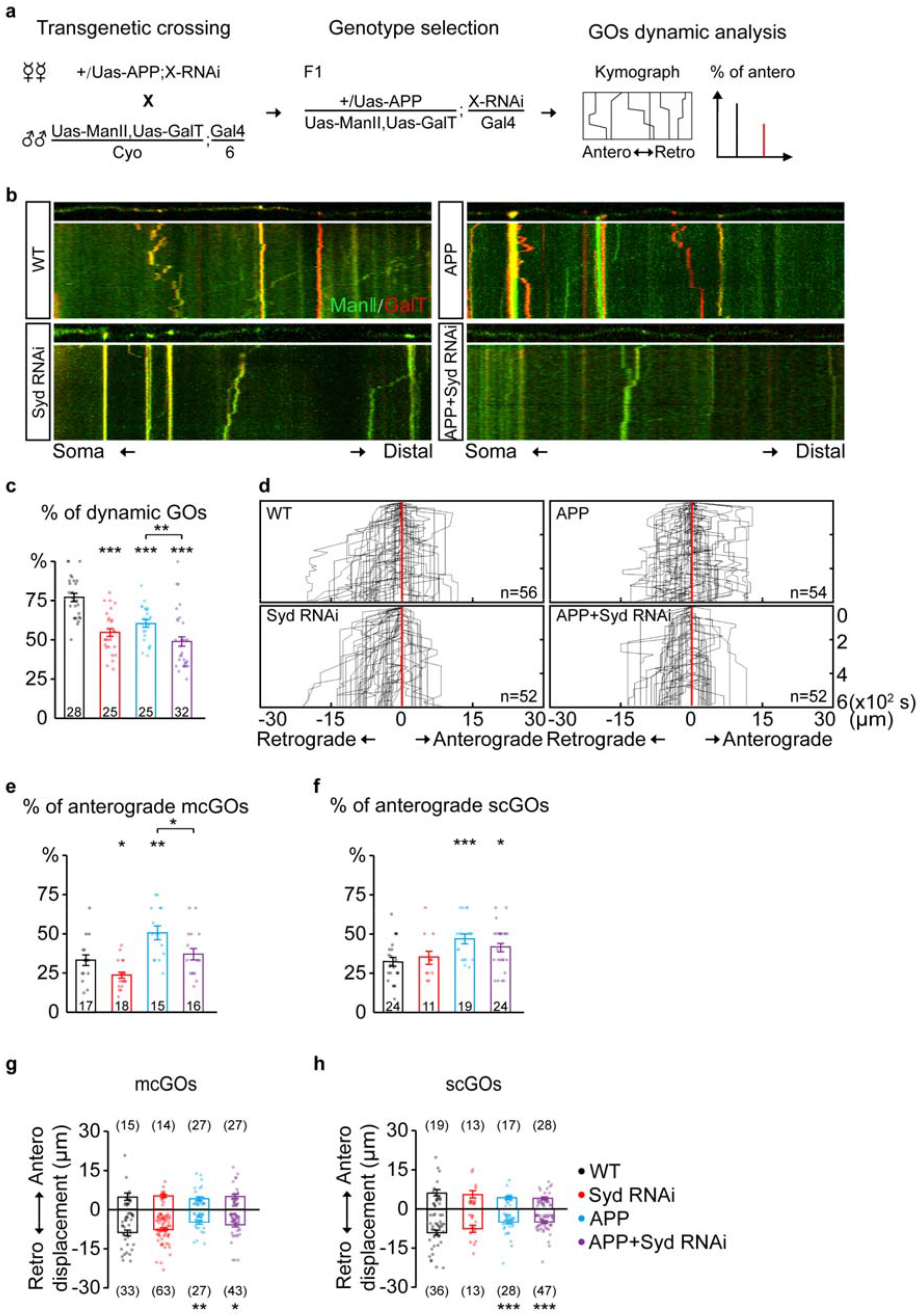
Syd is required for alteration of mcGOs dynamic induced by APP. **(a)** Flow of loss-of-function screening of candidate adaptor proteins which may be involved in GOs dynamic. **(b)** Examples of the trajectories of GOs movements in wild-type and APP neurons with Syd knockdown. Top: snapshots of the GOs in the straightened dendrites. Bottom: kymograph of time-lapse imaging of GOs in 10 min. Scale bar: 5 μm / 2 min. **(c)** Quantification of the proportion of dynamic GOs. **(d)** Compound tracks of GOs dynamic. **(e-f)** Quantification of the percentage of anterograde movements for **(e)** mcGOs, and **(f)** scGOs. **(g-h)** Quantification of the displacements for (**g**) mcGOs and (**g**) scGOs.

Further, we tested their function in APP neurons by examining three characteristics of GOs movements, including the motility of GOs, the direction and the displacements of dynamic GOs. The results showed that, for mcGOs, *Syd* RNAi restored the percentage of anterograde movements in APP neurons to normal (Fig 4e), while the loss-of-function of Lva and NudE did not (Fig S3b). Meanwhile, the motility of GOs and the displacements of mcGOs movements, as well as all the characteristics of scGOs movements cannot be recovered (Figs 4c and f-h, Fig S3c). These results suggest that the APP-induced alteration of the direction of mcGOs movements is Syd-specific, rather than Lva and NudE, although both Lva and NudE also are involved in regulation on the direction of mcGOs movements. Meanwhile, we checked the localization of Syd using fluorescent protein-tagged Syd (Syd-EGFP). The results showed that Syd present as puncta in C3da neurons, and colocalized with both somal Golgi and dendritic GOs (Figs S4a and b).

Taken together, our findings demonstrate that Syd specifically mediates the anterograde movements of mcGOs, and is also required for the regulation of APP on this process.

### Syd contributes to the defect of dendritic branching induced by APP

Dendritic defects are one of the pathological features of AD (S. J. Baloyannis, 2009). It has been reported that GOs are involved in dendrite development and morphogenesis (Horton et al., 2005; Ori-McKenney et al., 2012; Ye et al., 2007). Given the involvement of Syd in APP-induced abnormal GOs dynamic, the role of Syd in APP-induced dendritic defects was explored. We first characterized the dendritic morphology of C3da ddaA neurons with loss of Syd (Figs 5a-f). The results showed that Syd regulated both the features of dendrites and dendritic spikes. Compared with the wild-type, *Syd* RNAi led to an increase in the total number of dendritic branches, especially in high-order branch points (the fourth-order and up; Figs 5c and d), and a significant decrease in the density of dendritic spikes (Fig 5e). Meanwhile, the length of both dendrite and dendritic spikes showed no difference between *Syd* RNAi neurons and wild-type (Figs 5b and f). These results suggest that Syd is necessary for the maintenance of normal dendritic morphology.

**Fig 5.**
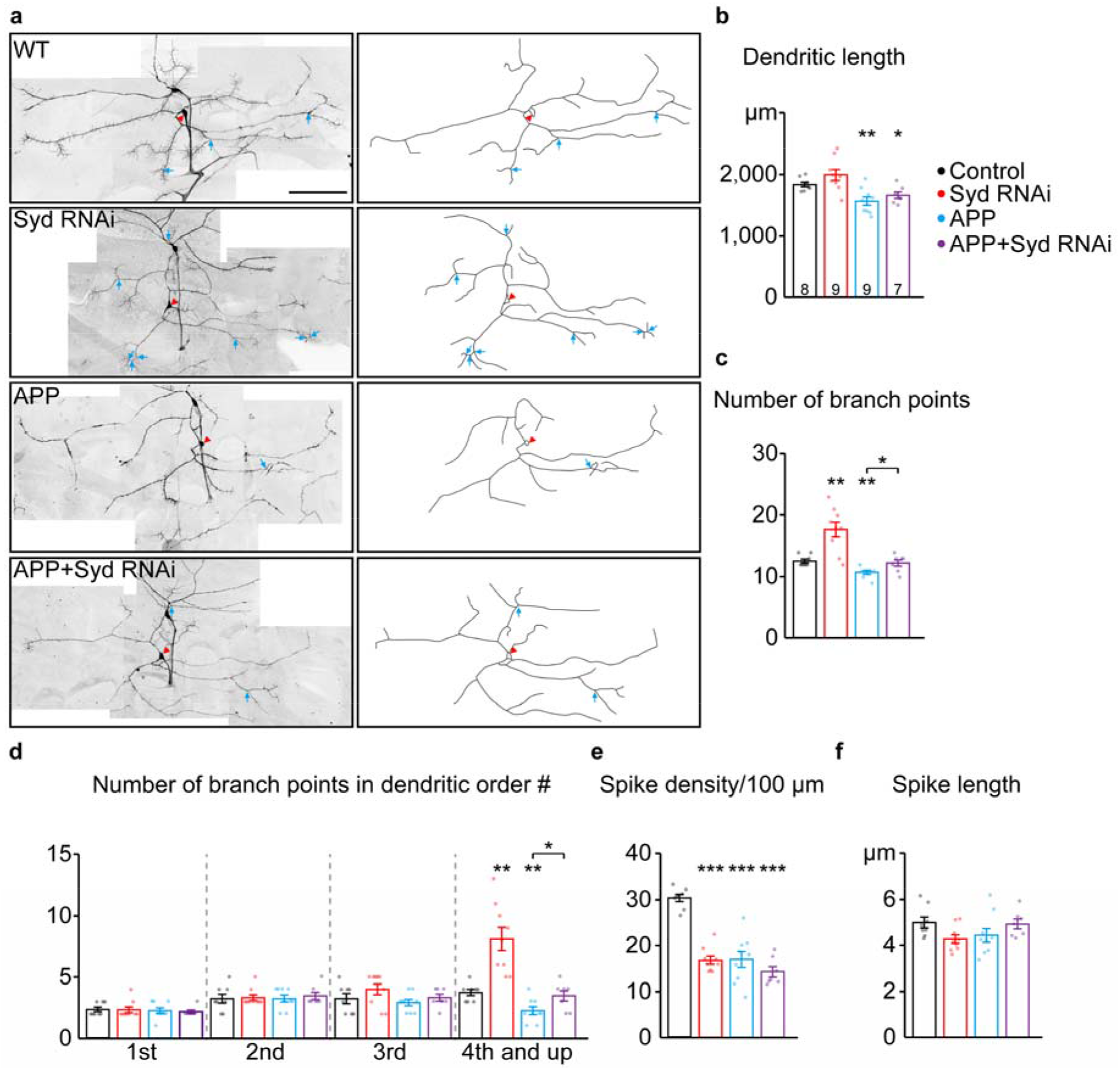
Loss of Syd recovers the dendritic defects induced by APP. **(a)** The dendritic morphologies of C3da ddaA neurons in wild-type and APP neurons following the loss of Syd. Raw images (left) and tracings of dendritic branches (right, the numerous F-actin-based “dendritic spikes” of C3da neurons are not included). The red arrowhead points to the soma and blue arrows to the branch points of the 4th order and up. Scale bar: 100 μm. **(b-f)** Quantification of dendritic morphologies. **(b)** The total dendrite length, **(c)** the number of total branch points, **(d)** the number of branch points in different orders; **(e-f)** The characteristics of dendritic spikes: **(e)** density / 100 μm, **(f)** average length.

Next, we examined the dendritic morphology of APP neurons after *Syd* knockdown. The results showed that the absence of Syd partially rescued the dendrite phenotypes in APP neurons but without affecting the characteristics of dendritic spikes. Consistent with the previously reported dendritic pathology in AD studies (Alpár et al., 2006; Spires et al., 2005), APP neurons showed a significant decrease in dendrite length, the total number of dendritic branch points, in particular high-order arbors, and the density of dendritic spikes (Figs 5b-e). *Syd* RNAi can restore the number of dendritic branches in APP neurons to normal, especially the number of high-order dendritic branches (Figs 5c and d). However, *Syd* RNAi failed to rescue the total dendrite length (Fig 5b) and spike density in APP neurons (Fig 5e).

Taken together, *Syd* RNAi and overexpressed APP have opposite regulatory effects on the complexity of dendritic arborization: *Syd* RNAi leads to an increase, while APP induces a decline. In particular, *Syd* RNAi can restore it to normal in APP neuron. For the density of dendritic spikes, their functions are the same: both of them cause a decrease; and Syd-RNAi cannot rescue it in APP neuron. And more, Syd don’t make any effort on the length of dendrites and dendritic spikes induced by APP. Thus, these results suggest that Syd specifically mediates APP-induced branching defects in dendrites.

## Discussion

Here, we revealed that the dynein subunit Dhc and adaptor protein Syd mediated the regulation of APP on the direction of mcGOs movements. First, we found that APP exhibited colocalization and co-trafficking with GOs. Especially, APP reversed the distribution of mcGOs in the dendrites. Next, knockdown tests showed that the anterograde and retrograde movements of mcGOs were driven by Dhc and Khc respectively, thus raising a model for the bidirectional movements of mcGOs. Further, Dhc rather Khc mediated the increased anterograde movements of mcGOs induced by APP. Finally, loss-of-function screening identified that Syd was required for APP to induce alteration of the direction of mcGOs movements. And, Syd also mediated the dendritic branching defect induced by APP.

The abnormality of the direction of GOs movements is thought to be a new feature for APP-caused Golgi defects. Previous studies have exhibited the correlation between APP and Golgi units in the dendrites. APP (Das et al., 2013) and its β-secretase BACE-1 (Das et al., 2016) colocalize with the interior of Golgi-derived vesicles in the dendrites, and the α-secretase ADAM10 traffics from GOs to synaptic membranes (Saraceno et al., 2014). Here, our results displayed the direct link between APP and GOs by showing their co-localization and co-trafficking in the dendrites. These indicate that GOs serve as the carriers of APP trafficking in the dendrites. Further, the impact of APP residency on GOs was discussed. We point out that alteration of the direction of mcGOs movements is an important feature for APP’s regulation on GOs, which leads to a reverse of the distribution of mcGOs, and then causes dendritic defects. These results indicate that APP not only impairs the somal Golgi apparatus, but also leads to defects of dendritic GOs, pointing to a new direction for Golgi defects in AD.

Our results reveal that Syd serves as a bridge for APP to alter mcGOs movements. Several molecules have been studied in regulation of GOs transport and distribution: dominant-negative Lva (Lva^DN^) changes the distribution of GOs (Ye et al., 2007); the localization of NudE at GOs inhibits the anterograde movements of GOs (Arthur, Yang, Abellaneda, & Wildonger, 2015). Here, we found a new regulator of GOs transport, Syd, which is the homologue of mammalian JIP3 in *Drosophila*. JIP3/Syd is involved in axonal organelles transport, such as lysosome (Gowrishankar, Wu, & Ferguson, 2017) and early endosome (Abe et al., 2009), and defects of JIP3-dependent axonal lysosome transport aggravate Aβ accumulation in AD model (Gowrishankar et al., 2017). Here we discover that Syd localizes at GOs in the dendrites. Furthermore, Syd specifically participates in the transport of mcGOs: *Syd* RNAi suppresses the anterograde movements of mcGOs but not scGOs. This demonstrates that, consistent with Dhc, Syd mediates the anterograde movements of mcGOs, suggesting that Syd is an adaptor protein between Dhc and mcGOs. Further, we discover that *Syd* RNAi restores change of the direction of mcGOs movements induced by APP, while neither Lva^DN^ nor *NudE* RNAi do. In summary, we propose that Syd, as the adaptor protein of mcGOs movements, specifically mediates APP’s regulation on the direction of mcGOs movements.

Our results also demonstrate the functional differences between mcGOs and scGOs. After two types of compartmental organization of GOs in dendrites are reported (Zhou et al., 2014), Valenzuela and Perez put forward a hypothesis that mcGOs and scGOs play different roles in the secretory pathway: mcGOs are responsible for long-distance transport of proteins to dendritic branch points and synaptic junctions, while scGOs deliver the proteins to the destination, thereby promoting local protein transport to the membrane (Valenzuela & Perez, 2015). Here, the results from the loss-function-of Dhc and Khc point out that the mechanisms between mcGOs and scGOs movements are different. Both Dhc and Khc can regulate the direction of mcGOs movements, while that of scGOs can only be regulated by Khc. Besides, this distinction is also confirmed by the fact that Syd only regulates the direction of mcGOs movements, but not scGOs. Moreover, we also find mcGOs rather scGOs play important roles in neuronal development: only mcGOs regulate dendritic branching through Syd.

GOs defects could be new features of AD. Dendritic pathology occurs in the early stages of neurodegenerative disorders (S. J. Baloyannis, 2009). Recently, dendritic pathology related to GOs has been studied in a variety of neurodegenerative diseases: Lrrk2 mutation, one of the pathogenic factors of Parkinson’s disease, suppresses dendrite arborization by regulating GOs dynamic (Lin et al., 2015); nuclear polyglutamine (polyQ), a putative factor of Machado-Joseph’s disease, induces defective terminal dendrite elongation by impairing GOs synthesis (Chung et al., 2017). In AD, we find that *Syd* RNAi not only restores the abnormality of the direction of mcGOs movements induced by APP, but also rescues the defect of dendritic branching. These imply that the abnormality of mcGOs dynamic contributes to dendritic defects, thereby suggesting that GOs defects can be considered as indicators for early stage of AD.

Meanwhile, this work also indicates that the regulation of APP on GOs dynamic is reflected in many aspects: APP alters the characteristics of both mcGOs and scGOs movements, including motility, direction and displacements. Here, we only reveal the regulatory mechanism on the direction of mcGOs movements. These results suggest that the regulation of APP on GOs movements is achieved through multiple routes. The mechanism underlying regulation on the motility and displacements of GOs movements requires further investigation, which will help to better understand the mechanism of GOs deficiencies in AD.

## Materials and methods

### Fly stocks and transgenic line

The stocks of flies published in this study included: GAL4^19–12^ (Xiang et al., 2010); UAS-GalT::TagRFPt, UAS-ManII::TagRFPt/GFP (Ye et al., 2007; Zhou et al., 2014); UAS-CD8::GFP (Lee & Luo, 2001); UAS-APP::GFP^CG^, UAS-APP695^N-myc^ and UAS-APP695-Swedish^N-myc^ (a gift from Lei Xue, Tongji University, China); the transgenic fly stocks for the motor proteins subunits and adaptor proteins were shown in table S1 and S2 respectively. To generate the UAS-Syd:: EGFP transgenic line, the *Syd* cDNA (#RE24340, BDGP) and *EGFP* cDNA were amplified by PCR and transferred into the vector pJFRC2-10×UAS-IVS-mCD8-GFP (plasmid #: 26214, Addgene, Cambridge, MA). The construct was then injected into embryos of PBac{y[+]-attP-3B}VK00033 to generate transgenic flies. All stocks and crosses were maintained in a 25°C incubator with 40-60% humidity.

### Dissection, immunostaining, and imaging

Third instar larvae were dissected in insect saline and fixed with 4% formaldehyde (PFA) for 40 minutes; permeabilized and washed in wash buffer (1X PBS containing 0.15% Triton X-100), and then blocked in 5% normal serum for 30 minutes. After washing, larval fillets were incubated overnight at 4°C with the primary antibody and 3 hours with the second antibody at room temperature, avoiding light. The antibodies included: chicken anti-GFP (1:5000, Aves Labs, Inc., Tigard, OR); Alexa647 anti-HRP (1:1000; Jackson ImmunoResearch Labs Inc., West Grove, PA); Alexa488 anti-chicken (1:250; Jackson ImmunoResearch Labs Inc., West Grove, PA). The larval fillets were then dehydrated by alcohol and transparentized with xylene before mounting on the slide with DPX (Sigma-Aldrich, St Louis, MO) for imaging. The C3da neurons in the dorsal cluster of the fourth to sixth abdominal segments of the larvae were imaged using a confocal microscope (FV1000, Olympus, Japan) under a 60x oil immersion objective (NA=1.42, Olympus, Japan). The images were captured at a resolution of 1024*1024 pixels and a z step of 0.4 μm in XYZ mode with sequential scanning. Images of whole neurons require multiple views to be spliced together.

### Morphological analysis

For dendritic morphology analyses, we manually traced the dendritic branches by the NeuronJ plugin (Meijering et al., 2004). The tracings were used to measure the dendrite length and count the number of dendritic branch points. The number and length of spikes were similarly obtained.

To quantify the colocalization between Golgi and APP (or Syd), we calculated the proportion of colocalized puncta. To count the number of total Golgi and total APP (or Syd) as well as the colocalized ones, the Golgi and APP puncta were separately cataloged in ROIs manager, and the puncta that overlapped with the puncta on the other channel were counted as a colocalization (as shown in yellow). For the purposes of analyzing the compartmental organization of GOs, we classified the colocalized GalT::TagRFPt and ManII::GFP puncta as mcGOs, while others as scGOs.

### *In vivo* imaging

*In vivo* imaging was performed as previously described (Zhou et al., 2014). Firstly, the 3^rd^ instar larvae were collected and anesthetized with ether. The orientation of larvae was adjusted until the ventral trachea was located in the middle of the view field, and then the larvae were mounted in a drop of halocarbon oil. The movements of larvae were further limited by pressing a coverslip mounted on top of vacuum grease. Time-lapse imaging of fluorescently tagged GOs in C3da neurons was acquired using a confocal microscope (FV1000, Olympus, Japan), 60x oil immersion objective. The settings for the two channels were: APP/ManII (488nm laser for GFP, emission filter: 500-525 nm) and GalT (543□nm laser for RFP, emission filter: BA560IF). Multichannel time-lapse images were captured at zoom 2 with a resolution of 512*512 pixels and collected in XYZT mode for 10 min at 6 s intervals.

### Quantification of GOs dynamic

To quantify the dynamic behaviors of GOs, we generated kymograph images (as X-T images shown) using ImageJ. The kymograph images were generated in four steps: stabilization, straightening, reslicing, and timing projection. First, images were stabilized using the Stabilizer plugin (Li, Feb. 2008.) to avoid fluctuation from larvae movements. We straightened the dendrites to accurately measure the distances of GOs movements in the dendrites. The dendrites beyond the first branch points were traced and straightened over the entire stack. The straightened images were resliced to reconstruct the corresponding slices across the time series. Finally, the resliced images were time projected to produce a kymograph of GOs in the dendrites.

Further, we determined the number of dynamic GOs and analyzed their properties, including direction, displacements. We compared the position of GOs at the start and end of the movement, and classified the movement as anterograde when the final position of GOs puncta relative to the starting point was towards dendritic ends, and retrograde when the final position of GOs was towards soma. The coordinates of GOs trajectories on the kymograph were measured to obtain displacement information during movements. Meanwhile, the mcGOs and scGOs were distinguished based on colocalization analysis for ManII and GalT. Finally, the GOs trajectories were traced and generated combined tracks of GOs by aligning the starting points of the tracings, as shown in Figs 2c, 3c, 4d. Besides, we defined the difference between dynamic and stationary GOs and classified them as stationary if they moved less than 0.5 μm in any direction, otherwise as dynamic. To circumvent the influence of the physiological state of neurons and the accuracy of calculation, only kymograph images with at least one moving GOs were analyzed.

### Statistical analysis

Data were analyzed using a two-tailed Student’s *t*-test throughout and p-value was used to determine whether a significant difference existed. Significance levels were presented as follows: *, P < 0.05; **, P < 0.01; ***, P < 0.001; n.s., no significance. Data were presented as mean ± SEM.

## Acknowledgements

We thank Lei Xue (Tongji University, China), Bing Ye (University of Michigan, Ann Arbor), and Yang Xiang (University of Massachusetts Medical School) for sharing fly stocks and the Optical Bioimaging Core Facility of WNLO-HUST for support with imaging systems. This work was supported by grants from the National Natural Science Foundation of China (No. 31871027), the Science Fund for Creative Research Group of China (No. 61721092), Fundamental Research Funds for the Central Universities/HUST (2016YXMS034, 2014TS015), the Director Fund of WNLO to W.Z. and the China Postdoctoral Science Foundation (No. 2015T80788) to J.C.

## Authors’ contributions

W.Z. and J.C. conceived this project. W.Z., J.C. and Q.D. designed the experiments. Q.D. conducted the experiments and analyzed the data. G.C. carried out molecular experiments. W.Z., J.C., Y.Z. and Q.D. wrote the paper.

## Supplementary

**Fig S1.**
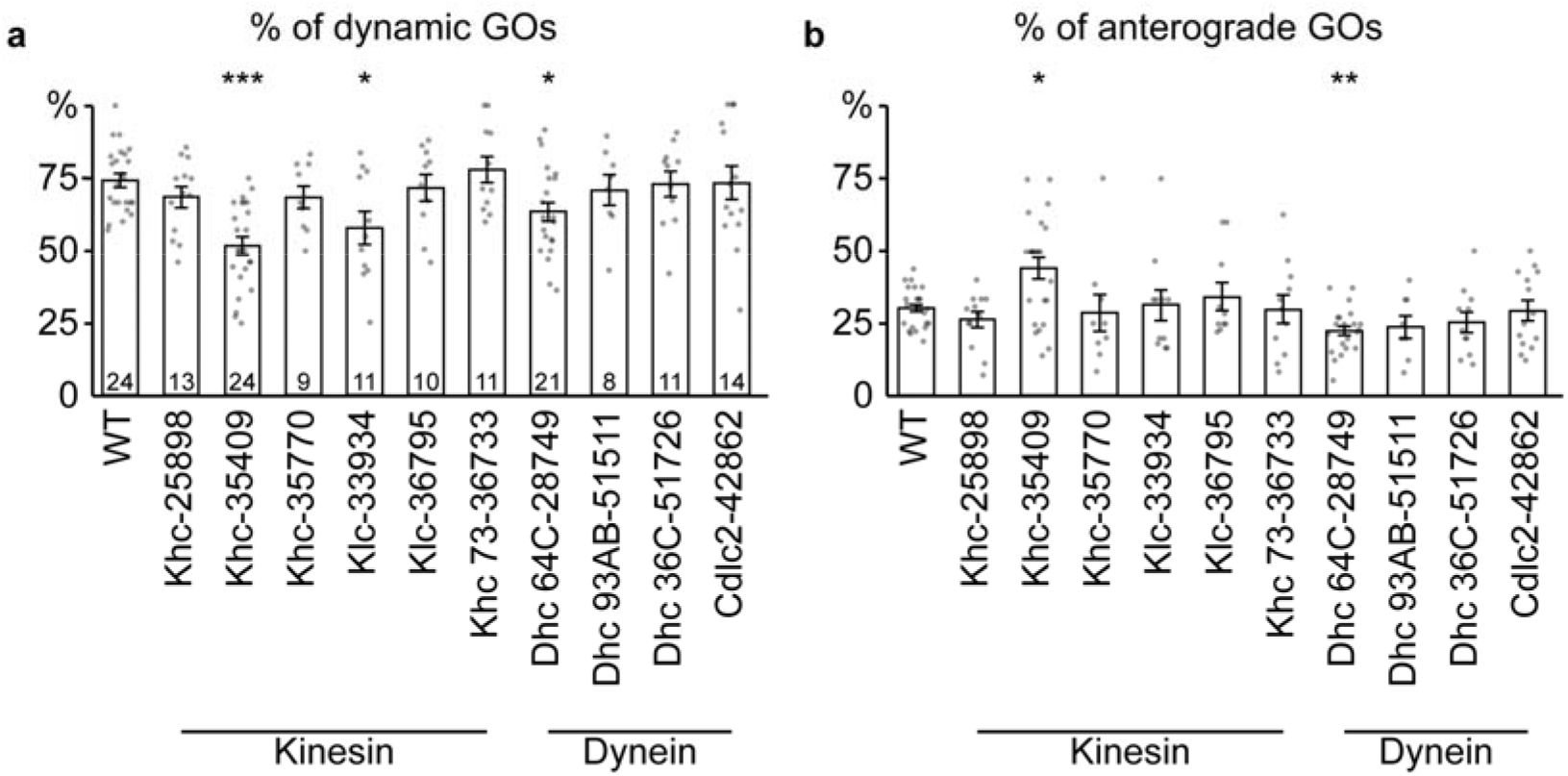
The RNAi-based knockdown tests identify motor protein subunits that regulate dynamic GOs. **(a)** The proportion of dynamic GOs, **(b)** the percentage of anterograde GOs movements.

**Fig S2.**
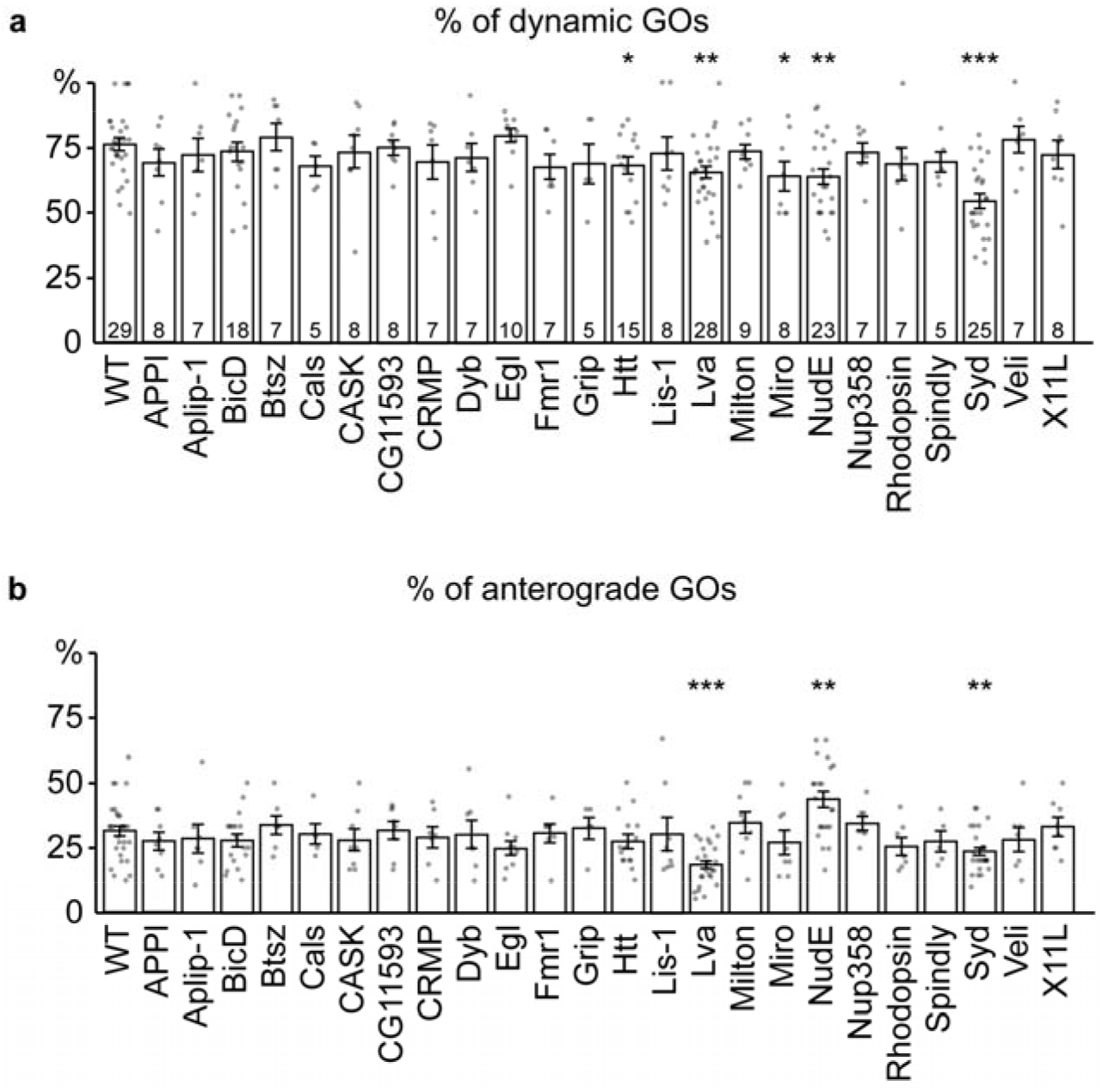
Screening results of the candidate adaptor proteins involved in the regulation of GOs dynamic. **(a)** Quantification of the proportion of dynamic GOs, **(b)** the percentage of anterograde GOs movements.

**Fig S3.**
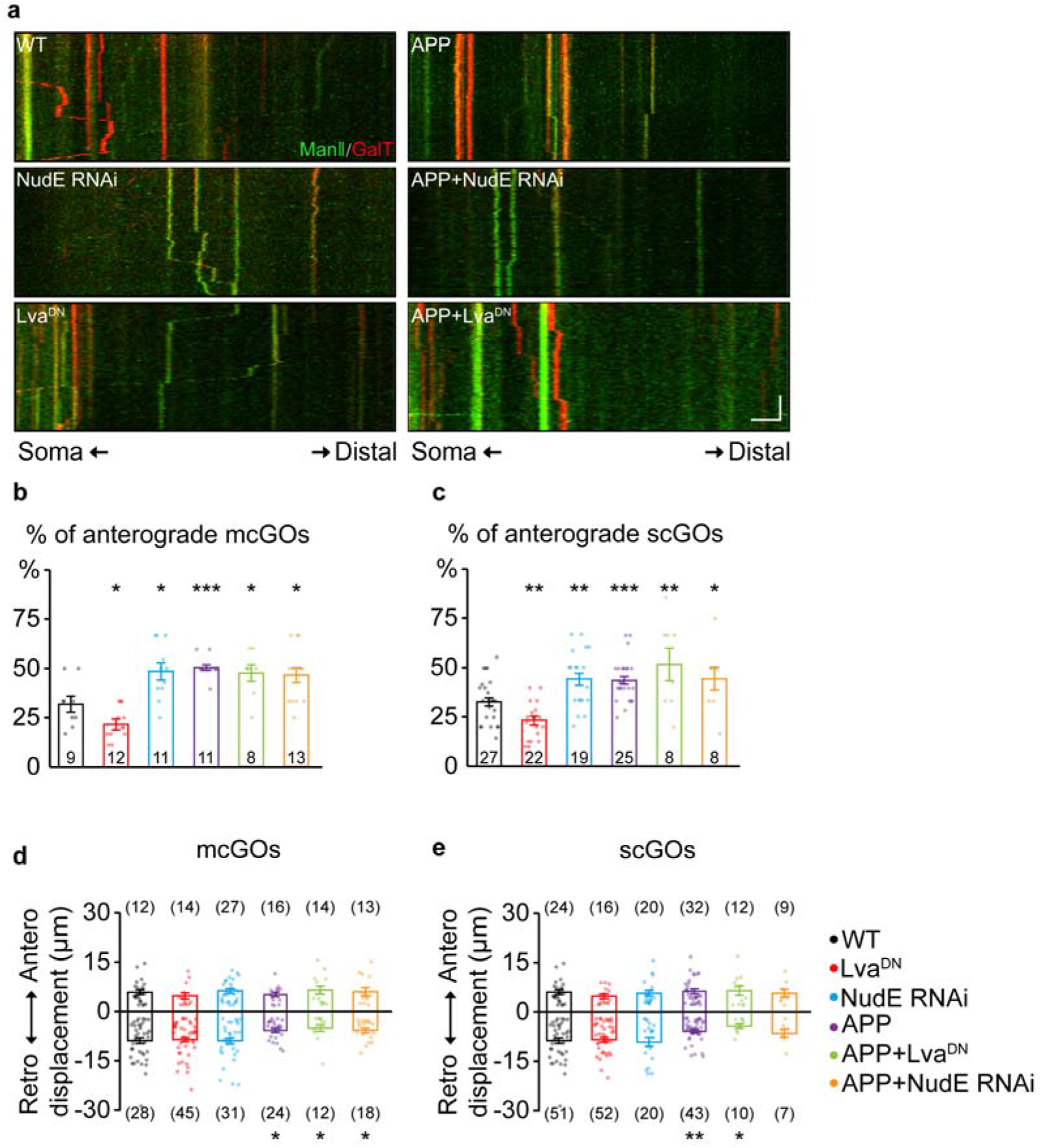
Neither Lva^DN^ nor NudE RNAi rescue the changes of GOs dynamic induced by APP. **(a)** kymograph showing the GOs movements in wild-type and APP neurons with Lva^DN^ or NudE-RNAi. Scale bar: 5 μm / 2 min. **(b-c)** Quantification of the percentage of anterograde movements for **(b)** mcGOs, and **(c)** scGOs. **(d-e)** Quantification of the displacements for **(d)** mcGOs, and **(e)** scGOs.

**Fig S4.**
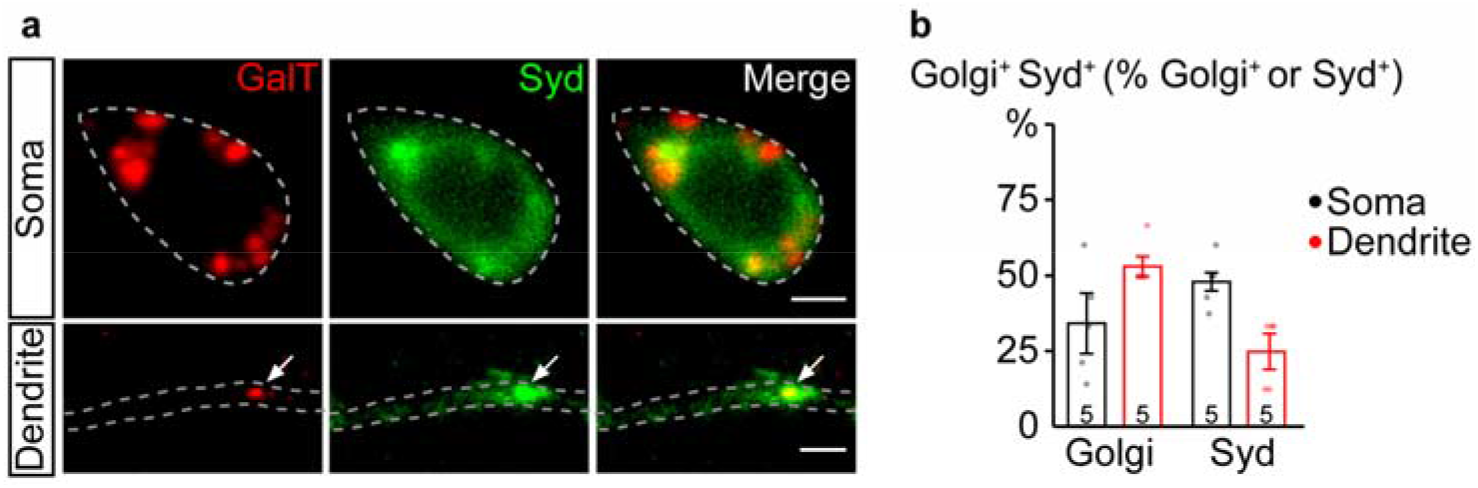
Syd localizes at GOs in dendrites. **(a)** Colocalization between Syd (green, Syd-GFP) with Golgi complex (red, GalT-TagRFPt) in soma and dendrite. Scale bar: 2 μm. **(b)** Quantification of colocalization between Syd and Golgi complex in soma and dendrites as the percentage of total Golgi complex (Golgi) or Syd.

**Table S1.**
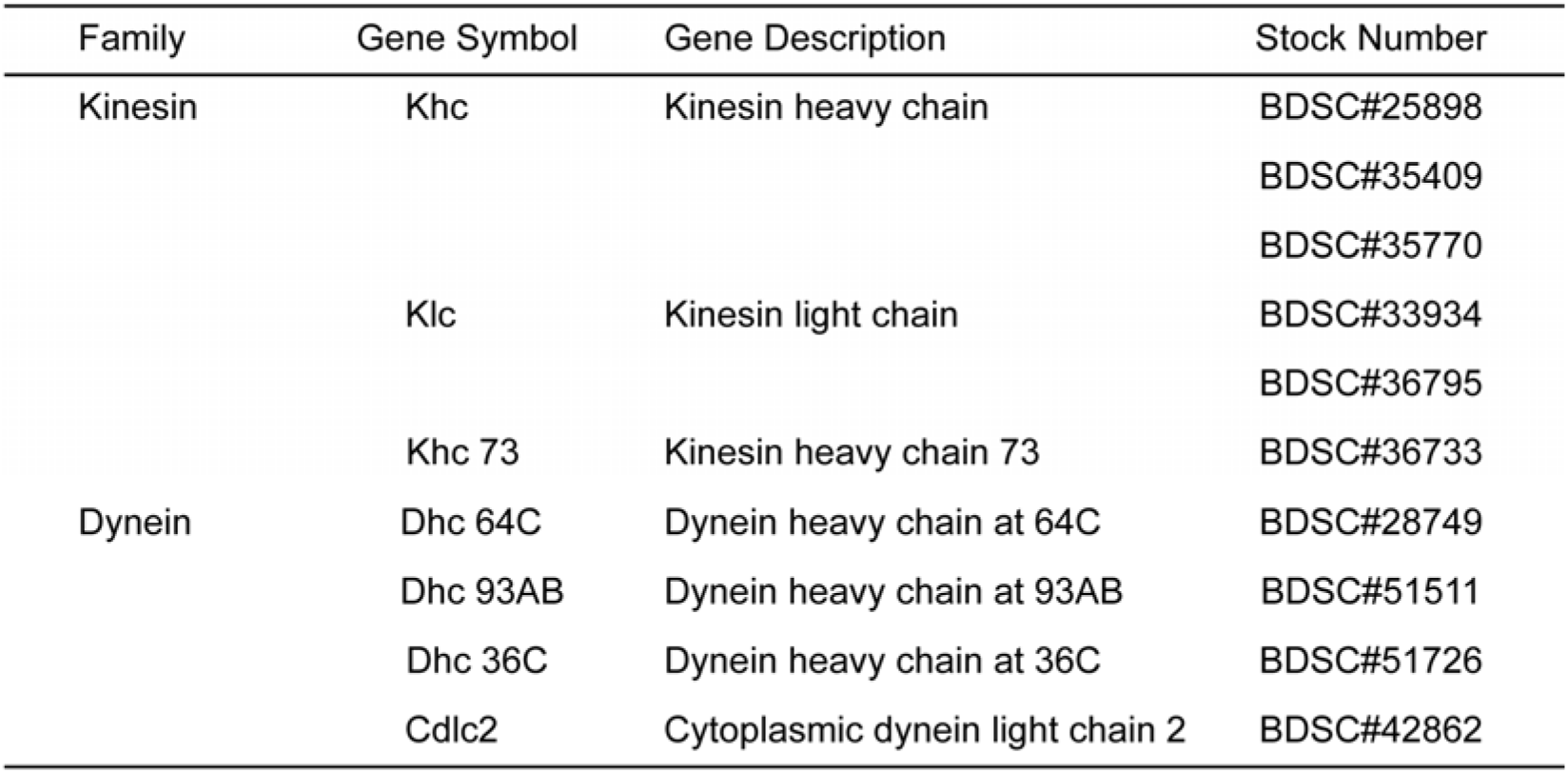
Trangenic RNAi fly stocks of the subunits of motor protiens

**Table S2.**
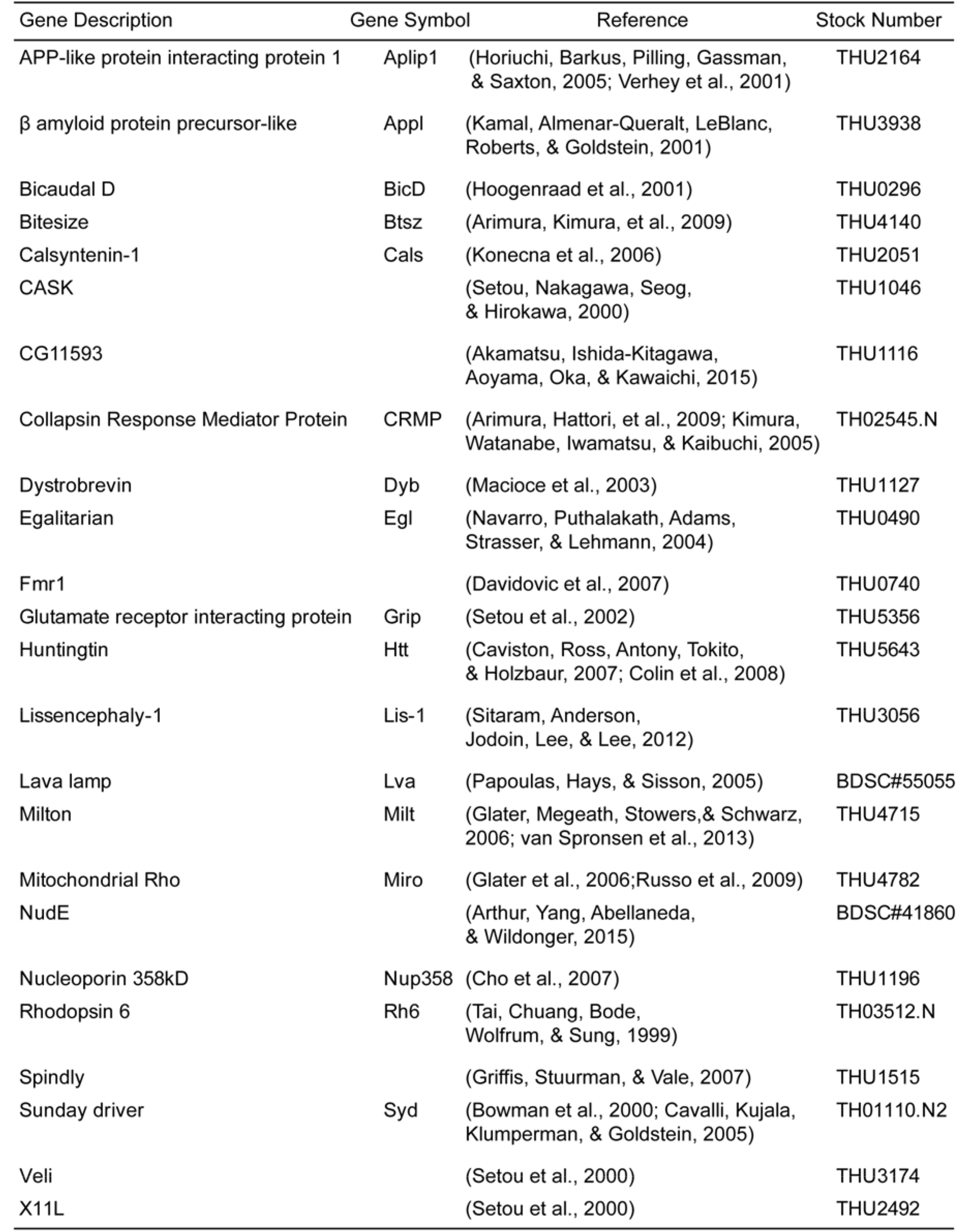
Trangenic fly stocks used for loss-of-function screening of adaptor proteins

| Gene Description | Gene Symbol | Reference | Stock Number |
| --- | --- | --- | --- |
| APP-like protein interacting protein 1 | Aplip1 | (Horiuchi, Barkus, Pilling, Gassman, & Saxton, 2005; Verhey et al., 2001) | THU2164 |
| $\beta$ amyloid protein precursor-like | Appl | (Kamal, Almenar-Queralt, LeBlanc, Roberts, & Goldstein, 2001) | THU3938 |
| Bicaudal D | BicD | (Hoogenraad et al., 2001) | THU0296 |
| Bitesize | Btsz | (Arimura, Kimura, et al., 2009) | THU4140 |
| Calsyntenin-1 | Cals | (Konecna et al., 2006) | THU2051 |
| CASK |  | (Setou, Nakagawa, Seog, & Hirokawa, 2000) | THU1046 |
| CG11593 |  | (Akamatsu, Ishida-Kitagawa, Aoyama, Oka, & Kawaichi, 2015) | THU1116 |
| Collapsin Response Mediator Protein | CRMP | (Arimura, Hattori, et al., 2009; Kimura, Watanabe, Iwamatsu, & Kaibuchi, 2005) | TH02545.N |
| Dystrobrevin | Dyb | (Macioce et al., 2003) | THU1127 |
| Egalitarian | Egl | (Navarro, Puthalakath, Adams, Strasser, & Lehmann, 2004) | THU0490 |
| Fmr1 |  | (Davidovic et al., 2007) | THU0740 |
| Glutamate receptor interacting protein | Grip | (Setou et al., 2002) | THU5356 |
| Huntingtin | Htt | (Caviston, Ross, Antony, Tokito, & Holzbaur, 2007; Colin et al., 2008) | THU5643 |
| Lissencephaly-1 | Lis-1 | (Sitaram, Anderson, Jodoin, Lee, & Lee, 2012) | THU3056 |
| Lava lamp | Lva | (Papoulas, Hays, & Sisson, 2005) | BDSC#55055 |
| Milton | Milt | (Glater, Megeath, Stowers, & Schwarz, 2006; van Spronsen et al., 2013) | THU4715 |
| Mitochondrial Rho | Miro | (Glater et al., 2006; Russo et al., 2009) | THU4782 |
| NudE |  | (Arthur, Yang, Abellaneda, & Wildonger, 2015) | BDSC#41860 |
| Nucleoporin 358kD | Nup358 | (Cho et al., 2007) | THU1196 |
| Rhodopsin 6 | Rh6 | (Tai, Chuang, Bode, Wolfrum, & Sung, 1999) | TH03512.N |
| Spindly |  | (Griffis, Stuurman, & Vale, 2007) | THU1515 |
| Sunday driver | Syd | (Bowman et al., 2000; Cavalli, Kujala, Klumperman, & Goldstein, 2005) | TH01110.N2 |
| Veli |  | (Setou et al., 2000) | THU3174 |
| X11L |  | (Setou et al., 2000) | THU2492 |

